# Ring attractor dynamics emerge from a spiking model of the entire protocerebral bridge

**DOI:** 10.1101/081240

**Authors:** Kyobi S. Kakaria, Benjamin de Bivort

**Affiliations:** Center for Brain Science and Department of Organismic and Evolutionary Biology, Harvard University, Cambridge, MA 02138.

**Keywords:** ring attractor, head direction, navigation, central complex, protocerebral bridge, ellipsoid body, leaky-integrate-and-fire model, bump

## Abstract

Animal navigation is accomplished by a combination of landmark-following and dead reckoning based on estimates of self motion. Both of these approaches require the encoding of heading information, which can be represented as an allocentric or egocentric azimuthal angle. Recently, Ca^2+^ correlates of landmark position and heading direction, in egocentric coordinates, were observed in the ellipsoid body (EB), a ring-shaped processing unit in the fly central complex (Seelig and Jayaraman, 2015). These correlates displayed key dynamics of so-called ring attractors, namely: 1) responsiveness to the position of external stimuli, 2) persistence in the absence of external stimuli, 3) locking onto a single external stimulus when presented with two competitors, 4) stochastically switching between competitors with low probability, and 5) sliding or jumping between positions when an external stimulus moves. We hypothesized that ring attractor-like activity in the EB arises from reciprocal neuronal connections to a related structure, the protocerebral bridge (PB). Using recent light-microscopy resolution catalogues of neuronal cell types in the PB (Wolff et al., 2015; Lin et al., 2013), we determined a connectivity matrix for the PB-EB circuit. When activity in this network was simulated using a leaky-integrate-and-fire model, we observed patterns of activity that closely resemble the reported Ca^2+^ phenomena. All qualitative ring attractor behaviors were recapitulated in our model, allowing us to predict failure modes of the PB ring attractor and the circuit dynamic phenotypes of thermogenetic or optogenetic manipulations. Ring attractor dynamics emerged under a wide variety of parameter configurations, even including non-spiking leaky-integrator implementations. This suggests that the ring-attractor computation is a robust output of this circuit, apparently arising from its high-level network properties (topological configuration, local excitation and long-range inhibition) rather than biological nitty gritty.

## Introduction

An animal navigating in its environment relies on landmarks to estimate its orientation and position (Collett and Graham, 2004). In the absence of visual cues, many animals maintain a representation of their heading and position without landmarks by continuously tracking their own motion (Etienne and Jeffery, 2004). These representations are tuned by visual information but can be generated in the dark, without any visual feedback (Varga and Ritzmann, 2016; Taube, 2007; Seelig and Jayaraman, 2015), presumably by exploiting self-generated motion cues like efference copy (Kim et al., 2015). By integrating heading and distance traveled, an animal can estimate its current position (McNaughton et al., 2007). One of the key components of this computation is continuous tracking of heading. This requires the continuous tracking of variables in angular coordinates, a computation that can be accomplished by “ring attractor networks” (Solovyeva et al., 2016; Skaggs et al., 1995; Zhang, 1996).

In theoretical models of ring attractor networks, neighboring nodes connect to form a topological ring. The value of an angular variable is encoded in the radial position of a “bump” of neural activity within this ring. This bump arises through the combined dynamics of short range excitation and global or long range inhibition between nodes of the ring attractor network (Knierim and Zhang, 2012; Skaggs et al., 1995; Zhang, 1996). Asymmetric excitation of neighboring nodes causes the bump to move in that direction as its previous position is inhibited. In mammals, ring attractors are thought to explain the dynamics of the head direction (HD) cells, which are primarily found in the thalamus and cortical areas associated with the hippocampus (Taube, 2007). Each HD cell is tuned to a particular head orientation and the direction in which the cell fires maximally is referred to as its preferred direction. Different HD cells represent different allocentric directions and the motion of the bump through the network of functionally (though not physically) ring-shaped network of HD cells encodes head orientation. Studying dynamics in this circuit is difficult as these neurons are spread throughout relatively large areas in of the brain and not spatially organized according to their preferred directions, making simultaneous monitoring of their activity challenging.

In insects, it was recently shown that a physically ring-shaped network of neuronal connections (neuropil) may function as a ring attractor (Seelig and Jayaraman, 2015) within the midline-spanning central complex (CX), of *Drosophila melanogaster*. Specifically, the Ellipsoid Body (EB) and Protocerebral Bridge (PB), appear to contain a neural circuit implementing a ring attractor. The EB neuropil has a closed ring shape in dipteran insects but is split ventrally and therefore roughly linear or bean-shaped in all other insect groups (Strausfeld, 1976). Due to the evolutionary conservation of morphological cell types in the CX, it likely retains ring-shaped functional connections in all insects (Pfeiffer and Homberg, 2014). Furthermore, the linear EB structure has been shown to encode the angular position of the sun in locusts, a continuous variable in angular coordinates, suggesting ring-like function without closed ring shape (Heinze, 2014; Homberg et al., 2011; Heinze and Homberg, 2007). The compact size and physical ring shape of this neuropil uniquely facilitates the study of ring attractor dynamics in an complete and intact circuit that can be simultaneously imaged in an awake behaving animal (Seelig and Jayaraman, 2015).

In a closed-loop behavioral setup, Ca^2+^ activity in putative dendritic processes of one neuronal population within the EB was shown to encode relative angular position of a vertical stripe on a 2-D LED screen (Seelig and Jayaraman, 2015). Seelig and Jayaraman noted several features of their circuit that are typical of ring attractor networks (Haferlach et al., 2007; Knierim and Zhang, 2012; Arena et al., 2013). The Ca^2+^ activity in the E-PGs was localized in a single bump at any one time and this bump moved in response to the animal changing its heading. Furthermore, the bump exhibited spatial stability, when it did fade after the fly had been stationary for long periods of time: it typically reappeared in the same location when the fly moved, an indication of storage of the bump in physiological pathways other than Ca^2+^, e.g. perhaps subthreshold voltages. The bump locked onto a single stripe when two competitor stripes were presented and was observed to jump between identical stripes from time to time.

These neurons exhibiting these ring attractor-like dynamics connect two of the neuropil that make up the CX, tiling the EB with dendritic arbors and the PB with presynaptic boutons. They are called E-PGs (Ellipsoid Body-Protocerebral Bridge-Gall neurons, called PBG1–8.b-EBw.s-D/Vgall.b in Wolff et al (Wolff et al., 2015), EB.w.s or “wedge neurons” in Seelig and Jayaraman (Seelig and Jayaraman, 2015) and EIP in Lin et al (Lin et al., 2013)), denoting the flow of information within them from the EB to the PB and Gall (a secondary structure immediately outside the CX). The EB and PB are notable for their division into columnar segments, known as glomeruli in the PB and wedges/tiles (Wolff et al., 2015) in the EB. These computational units contain many different neural cell types beyond those shown by Seelig and Jayaraman to encode angular position. In the PB, these have been recently characterized at the level of morphology using single-cell stochastic labeling methods(Lin et al., 2013; Wolff et al., 2015). The resulting catalogue revealed that of the approximately 18 classes of neurons within the PB, only three reciprocally connected the EB and the PB.

We sought to test the hypothesis that PB neurons implement a ring attractor, based on their connectivity as enumerated in these these recent mapping papers. Using a leaky integrate and fire model and simple connectivity rules, derived from light-microscopy resolution neuronal morphologies, we have found that a simple model recapitulates the bump of Ca^2+^ activity and essentially all of the *in vivo* dynamics previously observed (Seelig and Jayaraman, 2015). Furthermore, we have found that this circuit is robust to variation in synaptic weights, behaving as a ring attractor under a wide variety of parameters, perhaps indicating that computing a ring attractor is the primary evolutionary function of the reciprocal connection between the EB and PB.

## Methods

Simulations were run in MATLAB 2015a and 2016a (The Mathworks, Natick MA USA) using custom scripts. All code to recapitulate these results is available at: http://lab.debivort.org/protocerebral-bridge-ring-attractor-model

The circuit network structure was coded from the data in Wolff et al. (Wolff et al., 2015) per the rules described in the Results section below. Leaky-integrate-and-fire dynamics were implemented using Euler’s method to evaluate the following equation, with *Δt* = 10^−4^s:

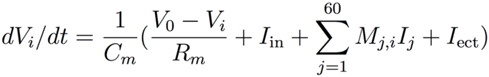

where *V*_*i*_ is the membrane voltage of neuron *i*, *I*_in_ is input current from neurons outside the PB circuit (0 in all neurons other than the E-PGs), *M*_*j,i*_ is the network connectivity matrix with entries equal to the synapse strength (in units of excitatory or inhibitory mini-postsynaptic currents (PSCs)), *I*_*j*_ is the output current of other neurons in the PB circuit, and *I*_*ect*_ is simulated ectopic current (such as might be induced by thermogenetic or optogenetic manipulation). We used parameter values that correspond to a generic spiking neuron, but these values are consistent with various *Drosophila* measurements or measurements of PB neurons in other species.*C*_*m*_ is the membrane capacitance (0.002µF in all neurons, assuming a surface area of 10^−3^cm^2^; (Gouwens and Wilson, 2009)), *V*_0_ is the resting potential (-52mV in all neurons; c.f. (Rohrbough and Broadie, 2002; Sheeba et al., 2008)), *R*_*m*_ is the membrane resistance (10MΩ in all neurons; (Gouwens and Wilson, 2009)), When a neuron’s voltage reached the firing threshold of −45mV (*V*_*thr*_; c.f. (Gouwens and Wilson, 2009; Sheeba et al., 2008)), a templated action potential trace was inserted into its voltage time series. This trace was defined as follows:

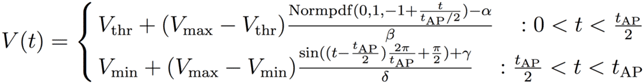

where *V*_*max*_ is the (purely cosmetic) peak action potential voltage (20mV; c.f. (Rohrbough and Broadie 2002)), *V*_*min*_ is the spike undershoot voltage (-72mV; c.f. (Nagel et al., 2015)), *t*_AP_ is the length of an action potential (2ms; c.f. (Gaudry et al., 2013; Gouwens and Wilson, 2009)), Normpdf (*a*,*b*,*c*) is the probability density function of a Gaussian with mean *a*, standard deviation *b* at *c*, and *α*, *β*, *γ* and *δ* are normalization parameters so that the max and min of the Normpdf and sin segments are 1 and 0 respectively prior to scaling by the voltage terms.

The firing of an action potential also triggered the addition of a templated postsynaptic current (PSC) trace to the output current time series of the firing neuron. The PSC trace was defined as follows in terms of *t* in ms:

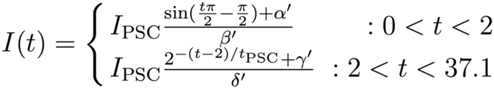

where *I*_PSC_ is the amplitude of a PSC (5nA; c.f. (Gaudry et al., 2013); excitatory and inhibitory PSCs were assumed to have the same magnitude but opposite sign), *t*_PSC_ is the time constant of PSC decay (5ms; c.f. (Gaudry et al., 2013)), and *α*’, *β*’, *γ*’ and *δ*’ are normalization parameters so that the max and min of the sin and exponential terms are 1 and 0 respectively prior to scaling by *I*_mini_.

Synapse strength parameters were explored manually to identify the baseline configuration in Figure 1. Thereafter parameter exploration was conducted as described in the Results. The overall magnitude of the synapse strength parameters shown in Figure 1 was the main free parameter of the model. The average synapse strengths of each synapse class are also free parameters, though we found that adjusting only the strengths of the Pintr>P-EG and Pintr>P-EN synapse classes was sufficient to recapitulate bump dynamics.

**Figure 1.**
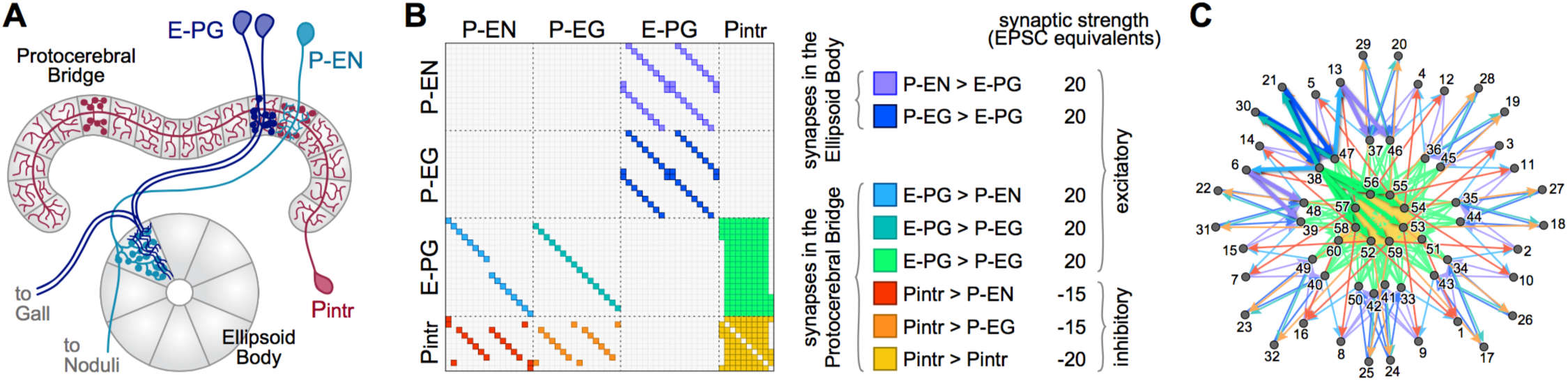
The Protocerebral Bridge neural circuit. **A**) Diagram of the PB and EB, illustrating three out of four modeled neural subtypes, the E-PGs, P-ENs and Pintrs. Not shown are the P-EGs which project from the PB to the EB. Axonal arbors are indicated with circular varicosities/ boutons. Dendritic arbors are intricate with fine linear branches. Overlap of an axonal arbor and a dendritic arbor within a single anatomical compartment (grey regions) is sufficient to postulate a synapse between neurons. Neurons with identical morphologies at the level of these anatomical compartments (e.g. the two dark blue E-PGs) are represented in the model as a single neuron. **B**) Matrix representation of the connectivity of the PB circuit. A filled rectangle in row *i*, column *j* indicates a synapse, with neuron *i* presynaptic, and neuron *j* postsynaptic. Different fill colors indicate different synapse classes, whose within-class strengths are drawn from a single distribution. In most implementations, the distributions of synapse strength associate with synapse classes have 0 variance, and means as shown at right. **C**) Graph, with node positions determined by a force-directed algorithm of the network with connectivity shown in B), which forms a ring with bilateral symmetry. Thick edges indicate lateral and reciprocal excitatory loops (local excitation) from neuron 38 (as an example) as well as excitatory connections to inhibitory neurons that target all glomeruli (long-range inhibition).

Leaky-integrator dynamics were implemented using Euler’s method to evaluate the following equation, with *Δt* = 10^−4^s:

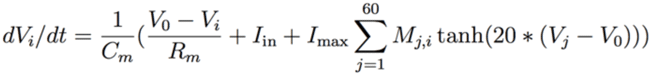

where all variables and constants are as defined above, and *I*_max_ is the maximum postsynaptic current achievable in a synapse of strength 1 within the PB circuit. First, the scaling parameter of the current-voltage tanh transfer function (20) was determined empirically. This value yielded dynamics that were the most bump-like, given the synapse strength parameters determined in the leaky-integrate-and-fire model. Then, synapse strength parameters that produced a fully functional bump were identified by adding Gaussian noise to the baseline parameters from the leaky-integrate-and-fire model. This noise had mean of zero and a standard deviation of 100% of the baseline value of each synapse parameter. The dynamics of approximately 200 such random configurations were examined manually, and those producing the best bump-like behavior were then iteratively refined using them as a new baseline, and then adding Gaussian noise with a standard deviation of 10% and then 5% of the respective baseline values. In order to break initial symmetry and allow the bump to move “spontaneously” random Gaussian noise with mean zero and standard deviation of 3x10^-10^V was added to each neuron in each time step.

Bump position was estimated and visualized by convolving the action potential rasters of each neuron with a Gaussian kernel with a standard deviation of 24ms. This approximates a Ca^2+^ signal in these neurons. Bump position was determined by taking the centroid, modulo eight, of this convolved representation for the P-ENs in each hemisphere. The estimated centroid of each hemisphere’s P-ENs was averaged to produce the final centroid estimate.

Circuit dynamics were captured for multidimensional analysis by simulating the network for 2 seconds, with inputs representing a rotating bar and two static competitors (setting parameters *SweepBarBool* and *TwinBarBool* equal to one in PBexperiment.m). 200 time points of the Gaussian-convolved spike rasters for each neuron were retained (from two sets of 2,000 contiguous frames each pulled from 200ms during the rotating bar and 200ms during the competitor bars, 200 time points were retained by decimating the data 20:1). Dynamics from 10,000 networks with randomly dithered synapse strength parameters constituted data points in this 200x60=12,000 dimensional space. To these the dynamics of networks in which a single synapse class parameter value was swept systematically from −9x to 10x its original value. The dynamics from these systematic sweeps were added to the dynamics from the randomly dithered networks and projected into two dimensions using PCA for visualization. Clusters of dynamics were enumerated using *k*-means clustering in the original 4,000 dimensional space. Representative dynamics of each cluster were computed by averaging all of the Gaussian-convolved spike rasters receiving each *k*-means cluster label.

## Results

To construct a circuit model of the PB we began with the catalogue of morphologically defined cell types in the PB (Wolff et al., 2015). This work enumerates all neuronal cell types within the PB, characterizing two cells as belonging to the same type if their pre- and postsynaptic arbors (as determined by MultiColor FlipOut imaging (Nern et al., 2015)) are in the same neuropil compartments. Compartments are defined as spatially distinct regions of the major glia-ensheathed neuropils of the central complex and associated regions. For example, the PB itself contains 18 glomerular compartments and the EB contains 16 wedge and 8 tile compartments. We included in our model 1) any neuron with postsynaptic processes in the PB and presynaptic processes in other compartments (output neurons), provided there is a PB input neuron with a postsynaptic arbor overlapping the presynaptic arbor of that output neuron, and 2) any neuron with presynaptic processes in the PB and postsynaptic arbors that overlap presynaptic arbors of neurons projecting out of the PB (input neurons; Figure 1). This includes all the neuronal cell types catalogued in Wolff et al. (2015) except for 5 classes of fan-shaped body projecting neurons (output only) and two classes of PB input neurons from the posterior slope (input only). We assumed that all neurons could be cleanly divided into dendritic and axonal compartments, and that information flows exclusively from the former to the latter.

The broad classes of neurons that met this criterion were the P-ENs (PB output neurons with axons in the EB and No), P-EGs (PB output neurons projecting to the EB and Gall), E-PGs (PB input neurons with dendrites in the EB and output to the Gall), and Pintrs (PB intrinsic neurons with both dendritic arbors and presynaptic boutons in the PB). P-ENs and P-EGs comprise 16 types each, defined by which PB glomerulus contains their dendrites. E-PGs comprise 18 types, defined by which PB glomerulus contains their axons (unlike “wedge” neurons (Seelig and Jayaraman, 2015) E-PGs also include neurons innervating the first and last glomeruli of the PB G9L and G9R (Wolff et al., 2015)). The Pintrs comprise 10 types, defined by which PB glomerulus contains their axons. If their projections were identical at the level of the 60 types described above, individual neurons were considered identical, and represented by a single neuron in the model. Lastly, we assumed that neurons formed no autapses. The connectivity of the network thus defined is shown in Figure 1B. We examined the topological arrangement of this network by using a force-directed algorithm (Fruchterman et al., 1990; Webb and Stone pers. comm.) to arrange nodes representing neurons. The connectivity present in this network has a ring-like topology, with bilateral symmetry (Figure 1C).

Circuit dynamics were implemented using leaky-integrate-and-fire (Stein, 1967) neuronal models, with values for the membrane capacitance, resistance, resting potential, undershoot potential, and postsynaptic current (PSCs) time constants and magnitudes, chosen to reflect generic neuronal properties. The important free parameters of the model were the strengths and signs of the synapses between each type of neuron. We assume that the strengths of all synapses between two classes of neurons (a “synapse class;” e.g. all synapses between P-ENs and E-PGs) were identical.

Strength of synapses was implemented as the number of PSC equivalents per action potential. Excitatory neurons induced positive, depolarizing currents in their postsynaptic partners and inhibitory neurons negative currents. We assumed all neurons were excitatory unless we had evidence otherwise. The Pintrs are glutamatergic (Gelfand et al., 2008) and possess connectivity similar to other inhibitory local neurons in spatially compartmentalized neuropils, e.g., the antennal lobe (Chou et al., 2010) and lateral horn (Fişek and Wilson, 2014), therefore we assumed they are inhibitory (Liu and Wilson, 2013).

To deliver inputs to the circuit, we assumed that information flows in first into the EB (this assumption has no bearing on our qualitative conclusions). Therefore, for each run of the model, the timing of action potentials in not-explicitly-simulated neurons upstream of the E-PGs was determined. These action potentials induced in the EBs excitatory currents with a strength equivalent of one PSC each. We assumed that background activity in these upstream neurons produced a Poisson-process sequence of action potentials with a mean rate of 5Hz. On top of this, Poisson-process spikes at higher rates (peaking at 120Hz) in subsets of E-PG types represented sensory-like input into the PB (Figure 2A), e.g. the azimuthal angle of light polarization (Heinze, 2014; Bockhorst and Homberg, 2015) or the retinotopic position of a landmark (Seelig and Jayaraman, 2013; Seelig and Jayaraman, 2015).

**Figure 2.**
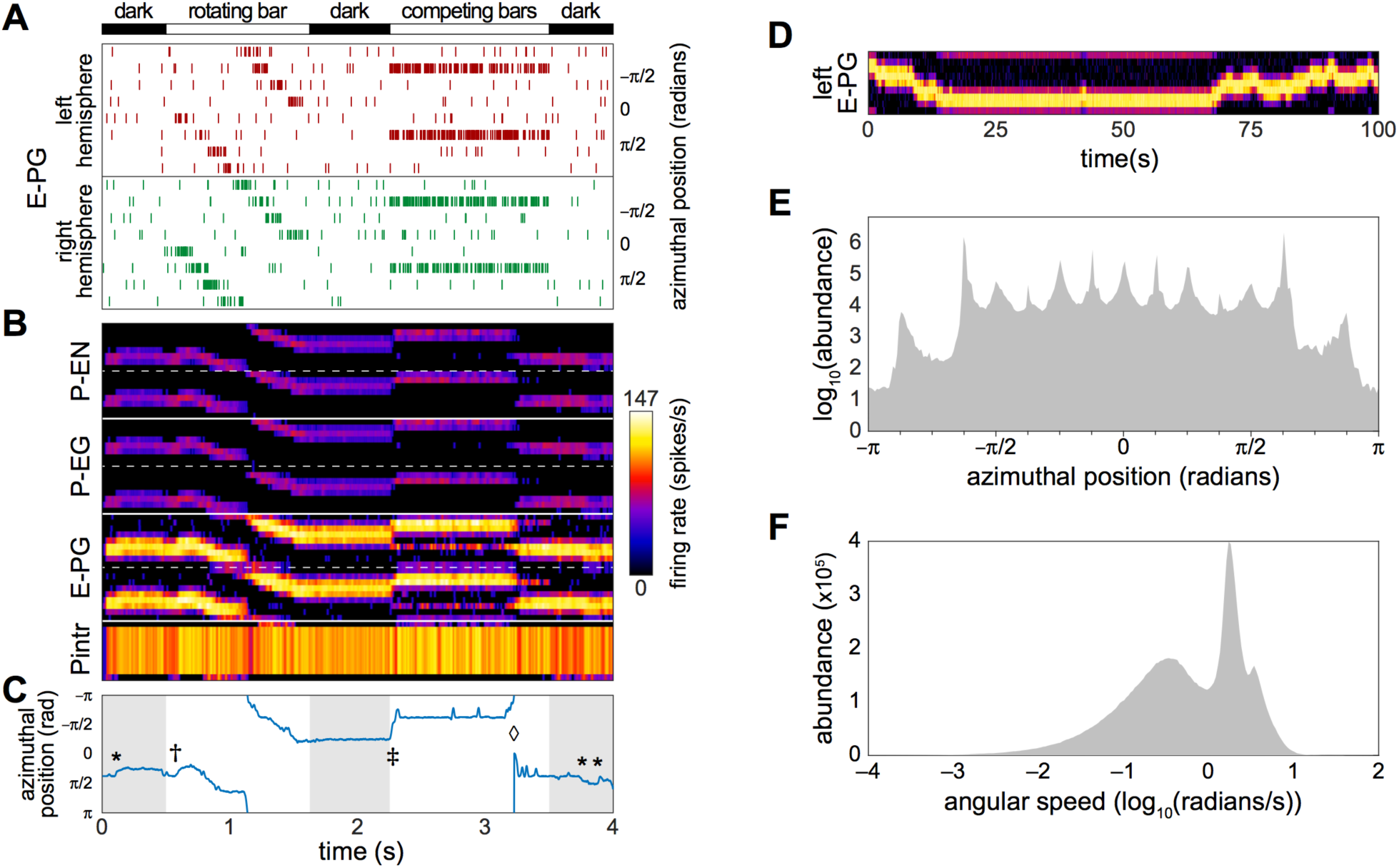
Bump-producing circuit dynamics. **A**) Input into the circuit is delivered as action potentials (raster marks) in neurons upstream of the E-PGs, plotted vs. time. Various sensory stimuli can be represented, including rotating bars, and static competitors. In “dark” periods, the only input is Poisson-distributed background activity. Inputs are associated with the 8 tiles(Tanya Wolff et al., 2015) of the EB, and corresponding azimuthal angles in body coordinates are indicated. **B**) Activity of all 60 neurons in the circuit versus time. Plotted heatmap is a Gaussian-convolved raster of action potential times (standard deviation 24ms). Dotted lines demarcate left from right hemispheres within a neural subtype. **C**) Position of the bump (centroid of activity in B; blue line) versus time. * indicates “spontaneous” shifts in the position of the bump in darkness. † indicates the bump sliding to the position of a bar as soon as it appears, and then following it as it rotates. ‡ indicates the bump jumping to the position of a single competitor when two static competitors appear. ◊ indicates the bump spontaneously switching its position to that of the other competitor. **D**) Bump behavior in darkness vs time over a longer period in the left hemisphere E-PGs. **E**) Histogram of bump frame-by-frame centroid position over 383 simulations of four seconds each in darkness. **F**) Histogram of spontaneous bump motion speed in the same dark simulations.

As a start of our characterization of circuit dynamics, we assumed, rather arbitrarily, that all synapse classes had a strength of 20 PSCs. With a small amount of manual parameter searching, we found that if the inhibitory synapses between the Pintrs and the P-ENs and those between the Pintrs and P-EGs had strengths of 15, circuit activity recapitulated several key phenomena that have been observed in Ca^2+^ recordings of the E-PGs (Figure 2B,C; Movie 1): 1) a stable “bump” of activity appeared at one position in the glomerular axis of the PB and the corresponding EB position, as observed by Seelig and Jayaraman (2015). This bump was almost always distributed over two or three glomeruli/tiles (25% to 38% of the azimuthal axis), corresponding roughly to the size of the Ca^2+^ bump they imaged (Seelig and Jayaraman, 2015). 2) The bump jumped or slid to the position of a novel sensory cue (i.e. a vertical bar), represented as increased firing rate in the neurons upstream of a single E-PG. 3) When the position of this input activity processed across adjacent glomeruli (moved in its azimuthal position), the bump followed. 4) When two competing vertical bars were provided in the form of firing-rate-matched activity upstream of two non-adjacent E-PGs, the bump moved to the position of one of the cues. 5) Occasionally, during the presentation of competitor bars, the bump would switch positions from one cue to the other. These characteristics were present for a wide range of synapse strength parameters (see stability analysis below).

As reported by Seelig and Jayaraman (2015), the bump appeared to be fairly stable in the dark (i.e. with only baseline background activity present upstream of the input neurons). Our baseline synapse strength parameter values yielded a bump “spontaneous drift rate” comparable to those observed *in vivo* (approximately 1 glomeruli/s; Figure 2D). We observed that the angular position encoded by the position of the bump had a highly discretized distribution while drifting in the dark (Figure 2E); the vast majority of the time, the bump was present in one of 15 azimuthal positions, and among these, ±5π/8 was the most abundant, followed by ±π/8. The distribution of bump speed during spontaneous motion (i.e. any motion in the dark) was trimodal (Figure 2F). These modes may correspond to staying in position, sliding between adjacent positions, and jumping between non-adjacent positions.

The emergence of a bump was remarkably robust, even a single action potential upstream of a single input E-PG was sufficient to induce a bump at the position of that action potential which would persist indefinitely (Figure 3A,B; Movie 1). We observed that occasionally the bump, as encoded by action potentials, would disappear briefly (up to a few tens of milliseconds at a time; Figure 3C-E). During these periods, none of the PB neurons would fire any action potentials, even if there were occasional action potentials in the neurons upstream of the E-PGs. This implies that the bump can be “stored” in sub-threshold potentials. These brief disappearances tended to happen when the bump was located at one of the less frequent azimuthal positions (e.g. ±7π/8).

**Figure 3.**
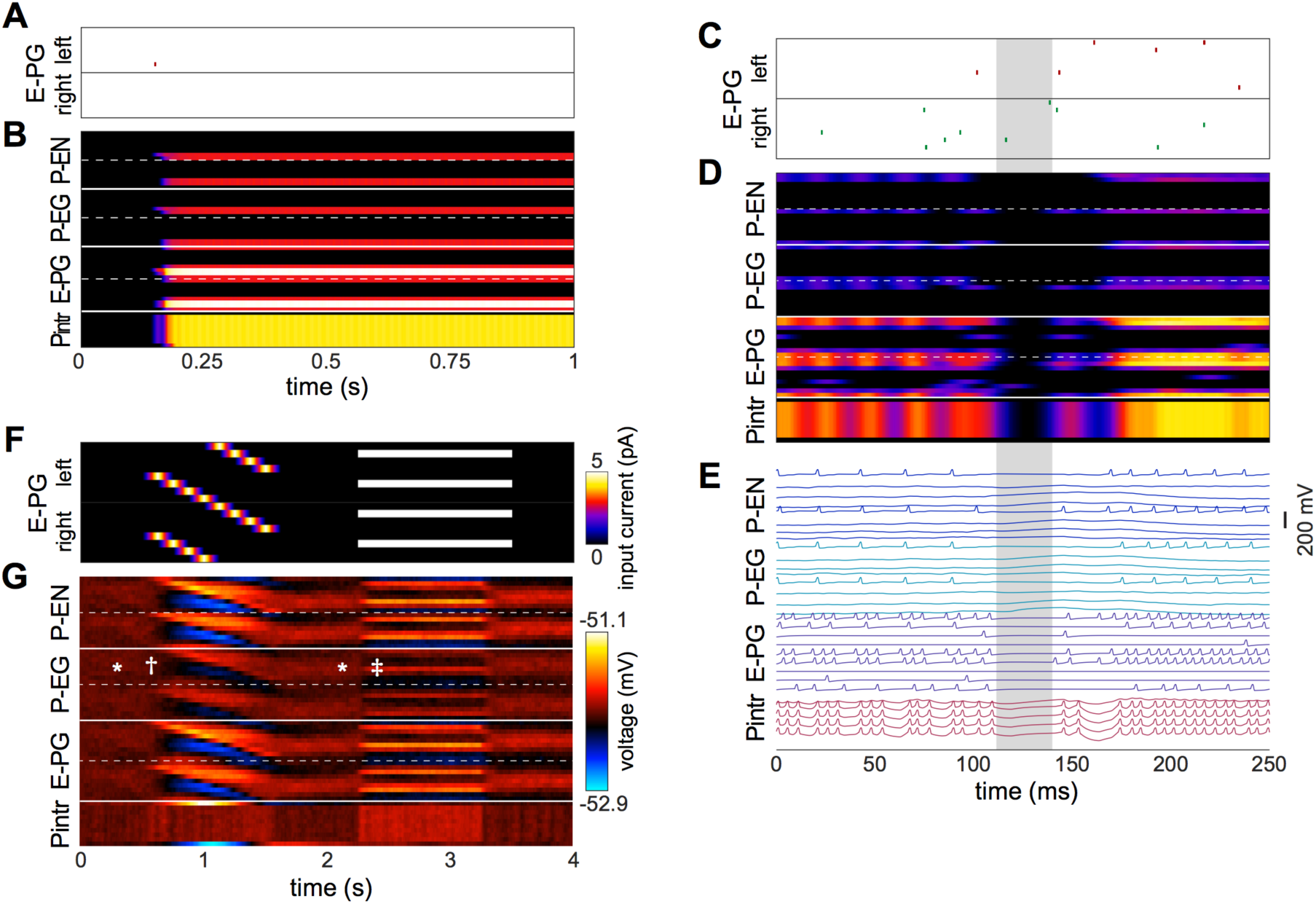
Relationships between the bump and action potentials. **A**, **B**) Input of a single action potential in an E-PG (red raster mark) is sufficient to induce a stable bump in the circuit. B) Gaussian-convolved raster of neural activity in all neural subtypes (axes and color scale as in Figure 2B). **C**) Circuit input via the E-PGs corresponding to 250ms of darkness. **D**) Corresponding dynamics of all neurons in the circuit revealing a ~20ms window (shaded grey) in which the bump disappears is not represented in action potentials, but reappears in the same position after the window (axes and color scale as in Figure 2B). **E**) Corresponding voltage trace. For clarity, the trace of every other neuron has been removed. **F**) Depolarizing currents representing input into the E-PGs in a leaky-integrator implementation of the circuit, versus time. Synapse strength parameters used were that provided dynamics most closely approximating a bump. **G**) Corresponding voltages in the entirety of the circuit. Symbols indicate elements of canonical bump phenomenology, as in Figure 2C. * indicates “spontaneous” shifts in the position of the bump in darkness. † indicates the bump jumping to the position of a bar as soon as it appears, and then following it as it rotates. ‡ indicates the bump jumping to the favoring one of two competitors (the lower competitor, most clearly discernible in the left hemisphere — the top half of each neuron type).

Several sets of neurons appeared to fire synchronously in the circuit (Figure 3E), specifically, those Pintrs that have axonal arbors in two PB glomeruli, bilaterally paired P-ENs and P-EGs, and bilaterally paired E-PGs (though this group of neurons is somewhat less synchronous by virtue of their being the input neurons that are stimulated at random times by upstream neurons). Leaky-integrator implementations (without action potentials) of this model could also produce a bump that persisted in the absence of sensory input, selected between competitor bars or formed a unitary bump after competitors were removed (Figure 3F,G). However, the bump in this implementation did not have the same rapid spontaneous bump formation, spatial precision, or strong selectivity between competitors seen in the leaky-integrate-and-fire implementation (though it did have weak selectivity between competitors).

We next examined whether we had lucked out in finding synapse strength parameters that recapitulated so many experimental bump phenomena. We added random, Gaussian-distributed noise (mean = 0, standard deviation = 20% of each parameter’s baseline value) to the synapse strength parameters and then stimulated these dithered circuits with inputs of 1) sequential bursts of activity in adjacent wedges representing a rotating bar and 2) elevated activity in two non-adjacent glomeruli representing stationary competitor bars (Figure 4A). For each of these configurations, the ensuing circuit activity in all neurons during diagnostic periods of this stimulation (200ms from the the rotating bar phase and 200ms from the beginning of the competitor bars) were treated as points in a high dimensional space of circuit behavior. These points were clustered and averaged within a cluster to provide an exhaustive catalogue of the modes of dynamics that this circuit topology can produce (Figure 4A,B). Of 15 modes, three feature sets of neurons with essentially no activity (modes 1-3) and five feature sets of seizing neurons (modes 11-15). Two of the remaining modes feature bumps exhibiting all the key properties observed experimentally (i.e. those shown in Figure 2; modes 4 and 5, which are distinguished largely by which competitor they select). Two modes have bumps that are stable on too-long timescales and extend over too many adjacent glomeruli, but otherwise show the key properties (modes 6 and 7). The remaining three modes (8-10) have some key bump properties, but are stable on too-long timescales, are too wide, and fail to select between competitor bars. Thus, it appears that bumps with the properties observed by Seelig and Jayaraman are a robust output of circuits with this topology under a wide range of synapse strength parameters.

**Figure 4.**
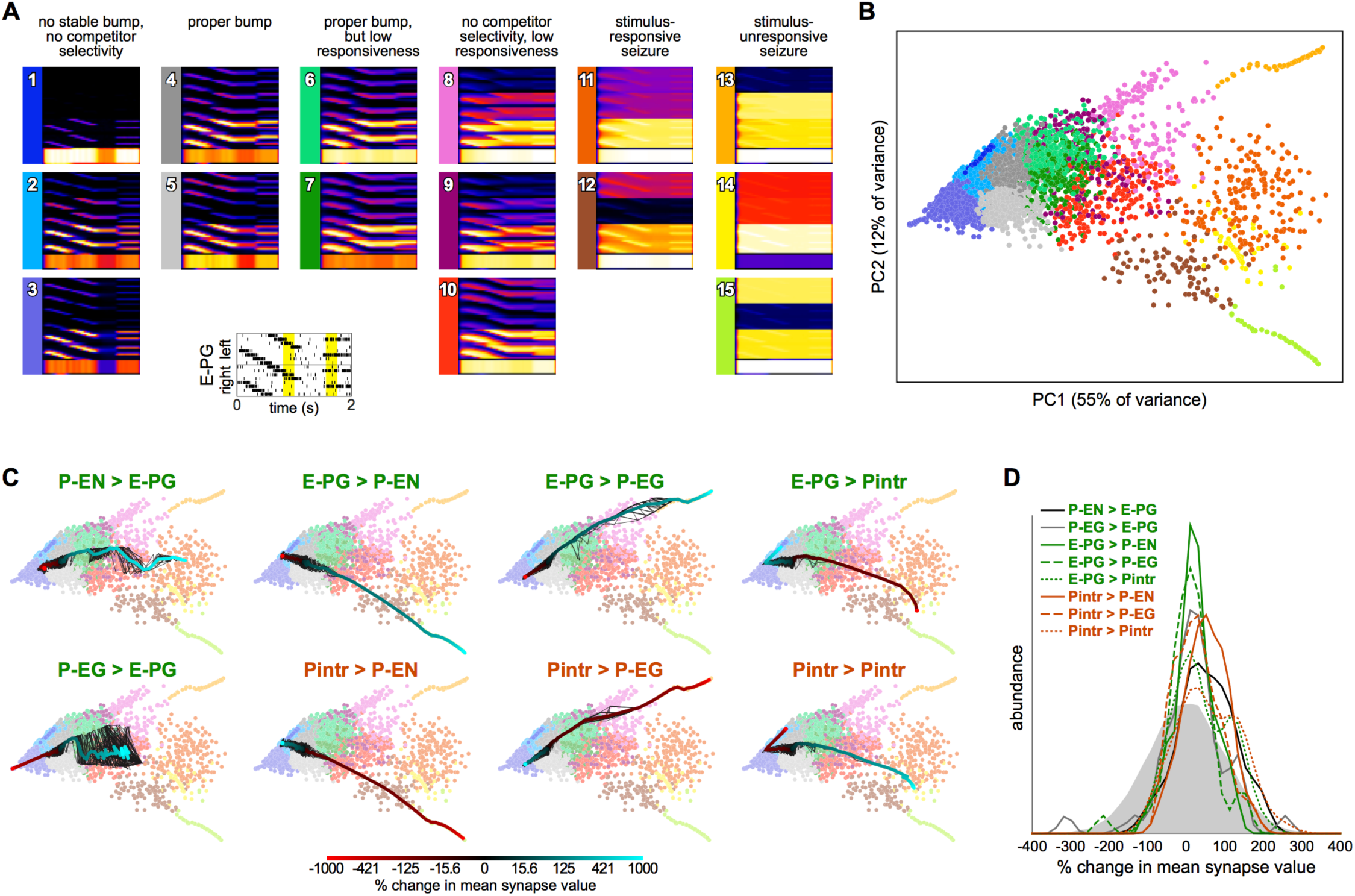
Robustness of bump dynamics. **A**) Modes of circuit dynamics. Each plot is an average of Gaussian-convolved action potentials rasters (as in Figure 2B). Constituent plots going into each average were identified by *k*-means clustering of circuit dynamics (B). For each synapse strength parameter configuration, the E-PG input shown in the inset was given. The resulting dynamics from the intervals marked in yellow were sub-sampled and clustered in 12,000 dimensions (200 timepoints for each of 60 neurons). Yellow regions are diagnostic of bump behavior since they include regions where the bump should follow a rotating bar and choose among competitors. **B**) Scatter plot of the first two principal components of circuit dynamics in the yellow intervals of A). Each point represents the dynamics of a circuit with synapse strength parameters equal to either 1) the baseline parameters (Figure 1B) plus Gaussian noise with mean = 0 and standard deviation = 20% of baseline value, or 2) the baseline parameters, with one parameter varied adjusted by −9x to 10x of its baseline value. Colors indicate 15 *k*-means clusters computed prior to PCA. Mean dynamics of all points within each cluster shown in A). **C**) Systematic variation of each of the eight synapse class strength parameter away from their respective baseline values. Black lines represent 10 different parameter value sweep replicates, and the thick color-mapped line their average, color coded by the shift of each respective parameter. **D**) Distributions of synapse parameters that support proper bump function. Gaussian noise from the solid grey distribution was sampled and added to each synapse class independently. This was repeated 24,000 times and the resulting circuit dynamics were k-means clustered into 400 clusters. 6 clusters were identified that had proper bump dynamics comparable to modes 4 and 5 of A). The distribution of offsets represented in these clusters is shown for each synapse class strength parameter.

To understand the contribution of each synapse class to circuit function, we systematically varied the strength of each synapse class from −9x to 10x its original value (Figure 4C). Converting excitatory drive from the PB to the EB into inhibition (by reversing the sign of either the P-EN>E-PG or P-EG>E-PG synapses) eliminated input-independent bump activity in the P-EGs (mode 3). Increasing the strength of that excitatory drive led to too-stable bumps without competitor selectivity (modes 9 and 10) and eventually seizure across the circuit (mode 11). Increasing inhibition of P-ENs (by either reversing the excitatory E-PG>P-EN synapses or amplifying the strength of the inhibitory Pintr>P-EN synapses), not surprisingly, eliminated activity in the P-ENs (mode 1). Conversely, the opposite manipulations resulted in a too-stable bump (mode 10) and eventually seizure of the P-ENs (and E-PGs and Pintrs; modes 12 and 15). Increasing inhibition of P-EGs (by either reversing the excitatory E-PG>P-EG synapses or amplifying the strength of the inhibitory Pintr>P-EG synapses), not surprisingly, eliminated input-independent bump activity in the P-EGs (mode 3). Conversely, the opposite manipulations resulted in a too-stable bump (mode 8) and eventually seizure of the P-EGs (and E-PGs and Pintrs; mode 13). Increasing inhibition of Pintrs (by either reversing the excitatory E-PG>Pintr synapses or amplifying the strength of the inhibitory Pintr>Pintr), resulted in too-stable bumps (mode 7), bumps with no competitor selectivity (mode 10) and eventually seizure in P-ENs, P-EGs, and E-PGs (mode 14). The opposite manipulations eliminated input-independent bump activity in the P-ENs (mode 2) and eventually all activity in P-ENs and P-EGs (mode 1).

This systematic variation of synapse class strength parameter values also provides evidence of the robustness of the bump phenomenon in this circuit. Increasing or decreasing the strength of a synapse class by up to 50% of its baseline value, for example, seldom changes the mode of circuit dynamics (Figure 4C). Thus, it seems a sizable parameter subspace around the baseline values can produce bump phenomena. This analysis allows us to assess how much of the parameter space around the baseline produces bumps, not how much of the total space can produce bumps. To discover more distant parameter configurations that might also work, we added a substantial amount of Gaussian noise to all parameters simultaneously (mean = 0, standard deviation = 100% of each parameter’s baseline value; this reverses the sign of a parameter 16% of the time). The vast majority of circuits with these more broadly-sampled random parameter configurations seized or were silent in at least one neural subtype but a small portion (~1.5% of 24,000 random parameter configurations) exhibited correct bump phenomena. These were identified and pooled by *k*-means clustering of circuit dynamics.

The distributions of each parameter within this pool of bump producing circuits are shown in Figure 4D. Each parameter can evidently take on a wide range of values, and with the right corresponding changes in other parameters, support bump function. Notably, almost all parameters could even have their sign reversed from excitatory to inhibitory or vice versa, and still contribute to a bump-producing circuit. The exceptions were the Pintr>P-EN and Pintr>P-EG synapses, which could be silenced but not converted into excitatory synapses, and still produce a bump. In general, however, the random noise that was added to the parameters in bump-producing circuits was positive, meaning that excitatory synapses could generally be made more excitatory, and inhibitory synapses more inhibitory, without loss of bump function. Several parameter distributions appeared to be multimodal, notably P-EG>E-PG, E-PG>P-EG, E-PG>Pintr, and Pintr>Pintr, suggesting there may be discrete (or non-linear manifolds of) synapse strength configurations that support bumps.

This framework allowed us to predict the effect of thermogenetic or optogenetic perturbation of neural populations. We computationally injected varying amounts of current into each neural subtype as defined by Wolff et al. (2015), i.e. distinguishing between E-PG and Pintr subtypes (Figure 5), and projected the ensuing circuit dynamics into the same space where we defined the dynamics modes (Figure 4B). In general the predicted effects matched the effects of changing the corresponding synapse class parameters. For example, injecting depolarizing current into the P-ENs had the same effect as increasing the strength of the excitatory E-PG>P-EN synapse class (or decreasing the strength of the inhibitory Pintr>P-EN synapse class). Injecting even relatively large (±5nA) currents into the gall-tip-projecting subset of E-PGs or the P_6-8_-P_9_ subset of Pintrs had little effect, presumably because these neural subtypes are less numerous in our model, represented by only 2 neurons each.

**Figure 5.**
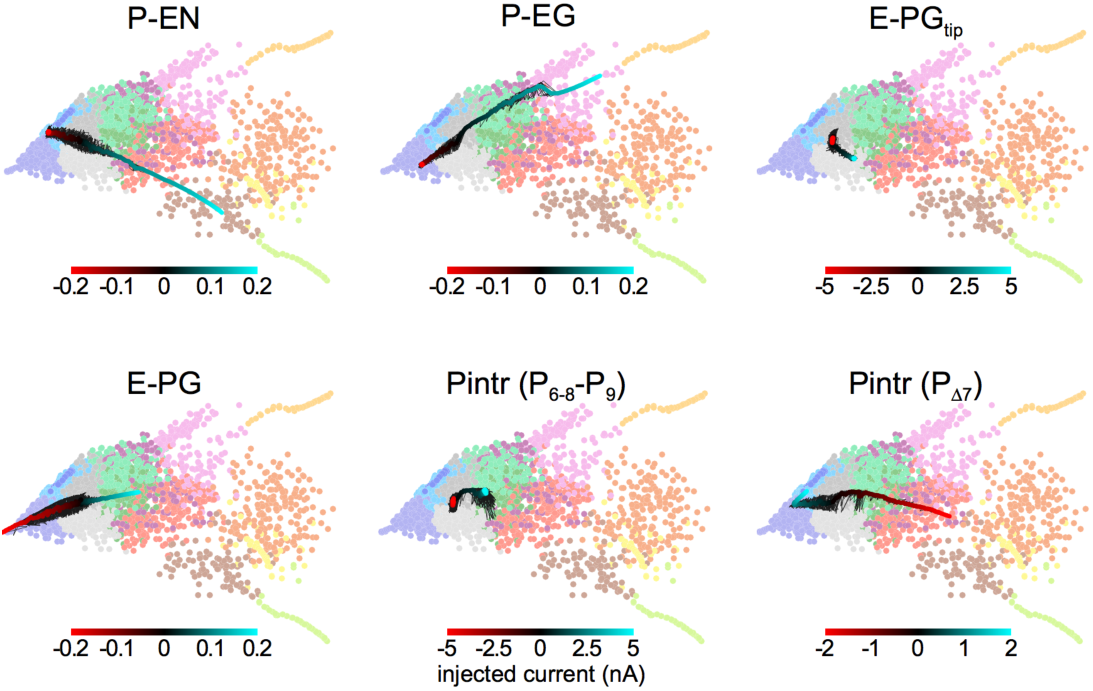
Prediction of circuit dynamics after neuronal subtype physiological manipulations. Circuit dynamics projected into two dimensions using the same input stimulus, diagnostic intervals, subsampling and linear projection as Figure 4B. Labeled neuronal subtypes were “injected” with ectopic currents as might be brought about by thermogenetic or optogenetic manipulation. Black lines represent 10 different current sweep replicates, and the thick color-mapped line their average. E-PG_tip_ refers to the subset of E-PGs that project to the Gall tip. Pintr(P_6-8_-P_9_) refers to intrinsic PB neurons that project from glomerulus 6-8 to glomerulus 9 and Pintr(P_Δ7_) that tile the PB with boutons while projecting dendrites throughout the PB.

## Discussion

Ring attractor networks are an attractive explanation for the storage and updating of continuous variables in the brain (Knierim and Zhang, 2012; Taube, 2007; Skaggs et al., 1995; Zhang, 1996) and may play a role in visual attention (de Bivort and van Swinderen, 2016). We have shown ring attractor dynamics arise in a network of generic spiking neurons with connectivity inferred from light-resolution microscopy and few other assumptions. The neurons in this network represent classes of neurons that are morphologically identical down to the level of independent computational units (glomeruli/wedges) defined in recent efforts to catalogue all neurons in the protocerebral bridge of the central complex (Wolff et al., 2015). The model produces a number of key behaviors that are predicted by ring attractor theory (Taube, 2007; Song and Wang, 2005) and observed by Seelig and Jayaraman (2015) in by Ca^2+^ activity in the E-PG neurons. In particular, a broad bump of activity (about 90-120° wide) tracks a simulated cue as it moves. We found that this bump may slide or jump to novel cues and chooses only a single cue if multiple competitors are provided (occasionally spontaneously jumping between them). Furthermore, we found that even when there is a pause in the representation of the bump by action potentials, it will reappear in the same position, as seen in Seelig and Jayaraman (2015). This suggests the bump is stored in subthreshold voltages.

Interestingly, our model suggests that there are discrete positions in the network in which this bump of activity prefers to reside as it moves through the network. Whether this is true of the circuit *in vivo* is not yet known, but it has been reported that startled cockroaches turn and run at angles that are multimodally distributed (Domenici et al., 2008). The modes of these escape angles are separated by approximately 30°, which is nearly matched by the 13 modes of bump position that we observed (Figure 2E). Perhaps discretized bump position tendencies underlie this distribution of escape angles. The distribution of bump speeds (Figure 2F) is also consistent with theoretical work predicting distinct types of bump motion: sliding between adjacent positions and jumping between non-adjacent positions (Zhang, 1996). Additionally, angular position vectors can be coded not only by which neurons are active, but also by which pairs of neurons have synchronous activity (Ratté et al., 2013). In our circuit we found that neurons tended to fire synchronously (Figure 3E), indicating that perhaps the PB could conceivably participate participates in a synchrony-based code.

We found that a large range of synaptic strength parameters can result in apparently proper ring attractor dynamics (Figure 4). Moderate levels of noise in the synaptic strengths within the circuit often still produced dynamics consistent with experimental observations. In our analysis we were able to characterize potential failure modes of the network, which include an inability to sustain the bump, low responsiveness, low or no competitor selectivity and/or network seizures. By systematically varying the synaptic weights of each class of connections, we explored the space of failure modes to evaluate the robustness of our model. Our model predicts that perturbing certain synaptic or neuronal classes could have larger impacts on this network than others. In general, neuronal classes with fewer neurons could be perturbed more dramatically before causing a breakdown in bump dynamics. At the same time, all the synaptic or neuronal classes could be dramatically perturbed (or even reversed in sign) and still produce a proper bump, provided appropriate compensatory changes in other classes were made. Going forward, our model may be able to provide a quantitative framework for understanding variability in individual differences in navigation, such as locomotor handedness (Ayroles et al., 2015; Buchanan et al., 2015).

It is important to consider which assumptions made in this model might not be realistic. The information flow of each neuron class is inferred from the overlap of “dendritic” and “axonal” cellular compartments determined by light-microscopy. Despite being unipolar, neurons in *Drosophila* generally have polarized information flow (Rolls, 2011), however, common axo-axonal, dendro-dendritic and perhaps even dendro-axonal synapses (Schneider-Mizell et al., 2016) paint a more complex picture. Electrical coupling, which can lead to synchronized neuronal firing, is also common in insect neurons (Pereda, 2014), but we have not included any in our model. Furthermore, we have assumed that every neuron has the same integration and firing dynamics despite the fact that the dynamics can vary significantly based on specific ion channel expression levels (Berger and Crook, 2015; Marder, 2011). We also make the assumption that if an axon and dendrite overlap in a compartment then they are connected, but this is not necessarily the case. Neurons that are adjacent with the resolution provided by light microscopy may not come into physical contact (Feinberg et al., 2008). Moreover, axons and dendrites which are in contact do not necessarily form functional synapses (Kasthuri et al., 2015). Due to these caveats, it is remarkable that our model recapitulates so many of the experimental observations of Ca^2+^ of E-PG neurons. The core computation of this circuit may be robust to many categories of biological detail, emerging instead from high-level connectivity of the sort that can be inferred from light microscopy.

Despite the conspicuous ring shape of the EB and its large number of inhibitory neurons with horizontal morphologies spanning all azimuthal positions (Martín-Peña et al., 2014; Kottler et al., unpublished), neither of these qualities is necessary to bring about ring attractor dynamics in our model. Instead, our model generates global inhibition using intrinsic PB neurons (the PBintrs; Figure 1). The EB has been shown to receive spatiotopic information about visual features from the bulbs (Seelig and Jayaraman, 2013) and is involved in visual place learning (Ofstad et al., 2011). These observations suggest that the EB encodes spatial information about landmarks in the environment which could be used to correct accumulated error in the position of a bump. While inhibitory circuitry within the EB is not required for ring attractor dynamics in the PB-EB circuit, we have no evidence that the inhibitory circuitry in the EB does not participate in a separate ring attractor. It is possible that both the Pintrs in the PB and the ring neurons (Martín-Peña et al., 2014) of the EB implement long-range inhibition for the production of two distinct ring attractors, which could interact to perform more sophisticated computations.

The egocentric heading correlate present in the PB-EB circuit is likely transmitted to other regions of the CX, particularly the fan-shaped body (FB). This neuropil could be a site of integration of navigational with internal state and sensory information for adaptive decision making. In addition to the PB-EB circuit neurons described here, the PB contains many columnar neurons projecting into the FB that have postsynaptic arbors in individual PB glomeruli and presynaptic boutons in different layers and columns of the FB (Wolff et al., 2015). Thus, it is likely that the FB inherits a bump or vertical band of activity from the PB. The FB is hypothesized to gate the selection of different behaviors in a state-dependent fashion (Weir and Dickinson, 2015) and activation of a single side of the FB induces ipsilateral turning (Guo and Ritzmann, 2013). Horizontal dopaminergic neurons in the FB have been shown to mediate sleep and arousal (Pimentel et al., 2016). The FB receives direct horizontal input from the visual system via the optic glomeruli (Ito et al., 2012) and also from many known modulatory neuropeptidergic neurons (Kahsai et al., 2012; Kahsai and Winther, 2011). The columnar projection neurons coming from the FB likely interact with these horizontal modulatory neurons. Therefore, it is appealing to hypothesize that the FB contains its own bump, downstream of the PB-EB bump, that it uses to integrate navigational information with neuromodulatory signals encoding internal states and sensory inputs.

## Author Contributions

BD conceived the project. KK and BD implemented the model, collected the data, analyzed it and wrote the manuscript.

## Acknowledgements

We thank Tanya Wolff and Vivek Jayaraman for helpful discussions and expert guidance on protocerebral bridge neurobiology. We acknowledge the National Science Foundation Graduate Research Fellowship Program and the Alfred P. Sloan Foundation for funding support.

